# Age-related behavioral resilience in smartphone touchscreen interaction dynamics

**DOI:** 10.1101/2024.03.01.583034

**Authors:** Enea Ceolini, K. Richard Ridderinkhof, Arko Ghosh

## Abstract

We experience a life that is full of ups and downs. The ability to bounce back after adverse life events such as the loss of a loved one or serious illness declines with age, and such isolated events can even trigger accelerated aging. How humans respond to common day-to-day perturbations is less clear. Here, we infer the aging status from smartphone behavior by using a decision tree regression model trained to accurately estimate the chronological age based on the dynamics of touchscreen interactions. Individuals (N = 280, 21 to 83 years of age) expressed smartphone behavior that appeared younger on certain days and older on other days through the observation period that lasted up to ∼4 years. We captured the essence of these fluctuations by leveraging the mathematical concept of critical transitions and tipping points in complex systems. In most individuals, we find one or more alternative stable aging states separated by tipping points. The older the individual, the lower the resilience to forces that push the behavior across the tipping point into an older state. Traditional accounts of aging based on sparse longitudinal data spanning decades suggest a gradual behavioral decline with age. Taken together with our current results, we propose that the gradual age-related changes are interleaved with more complex dynamics at shorter timescales where the same individual may navigate distinct behavioral aging states from one day to the next. Real-world behavioral data modeled as a complex system can transform how we view and study aging.

## Introduction

Aging is characterized by changes in brain and behavior. The age-related changes that slowly occur over the decades are typically described as a linear or curvilinear function of lived time (1–7). However, the trajectory of aging at shorter time scales may be way more complex (8). While some perturbations (e.g. disease) can accelerate aging, others (e.g. moderate exercise) decelerate or potentially even reverse it (9–13). According to the emerging study of ‘tipping points’ a complex system may have vulnerable periods when even a small perturbation can topple it into a distinct state (14, 15). Examples of such dynamics have been found in economics, climate sciences and ecology (15–18). To elaborate on one, an ecosystem can transition between lush green tropical forest, savannah and a barren tree-less state (19). The forest is vulnerable to climate change at tipping points where even a small change in climate (say reduced rainfall) may result in a catastrophic transition to savannah. Age-related brain and behavioral fluctuations may also occupy alternative stable states separated by tipping points (20).

The intrinsic ability to maintain brain function and behavior even under the influence of devastating perturbations such as in disease may play an important role in shaping the trajectory of aging (21). Nevertheless, this idea of *resilience* is clouded by distinct definitions and captured by subjective scales which mostly rely on the recollections of life events and how people cope or ‘bounce back’ from life stressors (22–24). The study of complex systems offers a generic and objective indicator of resilience based on how the system transitions between alternative stable states (16, 25). The underlying thinking is that the focus on isolated external perturbation is rather artificial, and instead the system should be considered as a whole under the continuous influence of interactive intrinsic and extrinsic stochastic forces (25). For systems with more than one stable state (basin of attraction), resilience is defined as the ability to absorb perturbations without being pushed into an alternative state (25, 26). This is mathematically realized by first fitting a model to the time series of fluctuations. The model captures the stochastic and the deterministic components. The model is then used to reveal the basins of attraction, and the resilience of the system can be estimated as the time taken to exit a basin and cross over the tipping point towards an alternative basin (25).

Modelling the essence of complex age-related dynamics requires a suitable marker that can reveal the fluctuations at short time scales. We draw inspiration from the ‘brain age’ marker based on neuroimaging (27, 28). In brief, the brain structure undergoes a range of complex age-related alterations, and machine learning algorithms can be trained to predict the chronological age (termed as *brain age*) using the rich brain images (27). In people with epilepsy or stroke, the gap between the *brain age* and chronological age increases by 6 to 10 years – as if the brain appears older in disease than in health (27). The model training leverages the inter-individual differences, but it can be repeatedly applied on the same individual to track slow intra-individual brain changes on the scale of months (10, 12).

We extend the approach of the *brain age* to smartphone behavioral data to derive ‘behavioral age’. To elaborate, smartphones are used in a broad range of behaviors – from dating to banking. One unobtrusive way to capture the rich behavioral repertoire is to focus on the touchscreen interactions. The inter-touch intervals (ITI) show a heavy tailed distribution, but this distribution is too low dimensional to capture the rich repertoire (29). The joint interval distribution (JID) offers a novel way to capture the various smartphone behaviors according to their next interval dynamics (30). In this distribution, the time interval between touchscreen interactions is considered in conjunction with the next interval k+1 (30). The ITIs are accumulated over a set period and captured in 2500 two-dimensional bins spanning the joint-intervals ranging from a few milliseconds to a minute (**Fig. 1A**). For instance, in the JID, the fast consecutive intervals underlying say typing a message are separately represented from the transition between fast to slow intervals say when switching from typing to reading. When the behavioral data is accumulated over hours to days, the JID contains information on behavioral processes, cognition and disease status (30–33). When accumulated over months, the inter-individual differences of the joint interval distribution are particularly strongly correlated to chronological age (33). A decision tree regression model can predict the age with a mean absolute error of 6 years by using the distribution as input (32), which is similar to the error range of 3 to 8 years achieved for *brain age* (27). Moreover, in people with epilepsy or stroke the gap between the *behavioral age* and chronological age increases by 6 to 10 years - as if the behavior appears older in disease than in health (32).

**Figure. 1.**
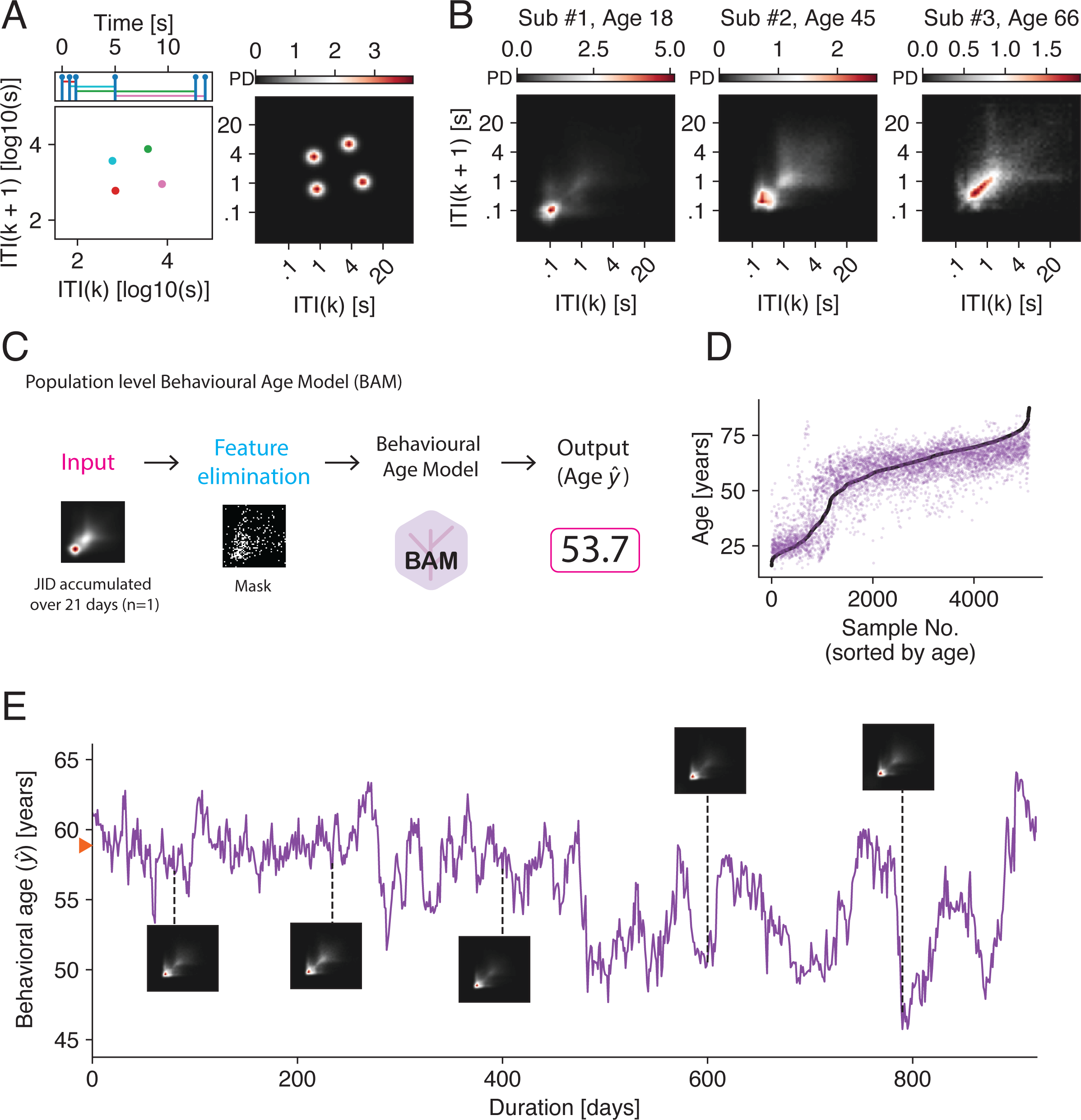
Behavioral age based on day-to-day smartphone touchscreen interactions. (A) We captured the diverse next-interval dynamics in smartphone interactions using the joint-interval distribution (JID). Scatter plot of a short simulated time series of next intervals to illustrate the method (left). The simulated events are on the top of the scatter plot. Note, how the short consecutive intervals (red) are separable from the slower next interval dynamics (green). Probability density in two-dimensional bins of the same scatter plot (right). Inter-touch interval, *ITI*. (B) The distributions accumulated over a period of 21 days showed substantial inter-individual differences. Here we show three different participants to illustrate the differences. (C) The behavioral age model (BAM) predicted the chronological age based on the smartphone JID – we termed these predictions as *behavioral age*. (D) The *behavioral age* of n = 5093 samples from N = 776 individuals vs. the chronological age. An individual contributed with one sample accumulated over a period of 21 days separated by 2 months. (E) The BAM was repeatedly applied on the same individual converting the intra-individual differences in the JID into a timeseries of *behavioral age*. The daily predictions were based on the JID accumulated over the previous 21 days. The model was trained and applied in folds such that the model trajectories were from individuals not seen by the model during training.

Akin to the *brain age* model, the *behavioral age* model when applied to a time series of smartphone behavior may capture longitudinal age-related fluctuations (34, 35). A key distinction is that the latter model can result in daily outputs to capture the day-to-day age-related behavioral fluctuations. It is not clear if the brain structure alters at the spatial resolution of the neuroimaging from day-to-day and even if it did the imaging method needs substantial resources rendering it infeasible to use daily (36). The smartphone next-interval dynamics on a given day is expected to differ from the previous day (30, 31, 37) and the *behavioral age* model allows the interpretation of these rich fluctuations in terms of appearing *older* or *younger* than the previous day. As the raw inputs to the model have negligible measurement noise (as in the behavioral events are always captured (38)) it is safe to assume that any fluctuation of the *behavioral age* stems from a true change in the next-interval dynamics. Here we trained a decision tree regression model to estimate the *behavioral age* from smartphone data gathered over 21 days. Next, we applied this trained model to obtain moving predictions resulting in the time series of *behavioral age* at the resolution of a day. Using this time series we address whether there are basins of attraction and tipping points in aging. Furthermore, we addressed whether behavioral resilience (absorbing the underlying forces of change without tipping into an alternative state) alters across the adult lifespan. Finally, we link the short-term dynamics on the scale of weeks to months to the slow fluctuations spanning decades.

## Results

### Time series of behavioral age based on the next-interval dynamics of smartphone interactions

Our analysis is based on individuals who identified as healthy at the time of recruitment (for subject selection see Fig. S1). We further confirmed their subjective health status using the SF-36 questionnaire (Fig. S2). We trained a population-level model to estimate the *behavioral age*. The model used data from N = 776 individuals, yielding a total of n = 5093 samples separated by 2 months (16 - 87 years of age). After the training, we applied the model daily at the level of each individual with greater than a year of continuous behavioral recordings (N = 291 individuals). The amount of smartphone usage was only marginally correlated with the chronological age (*β*_age_ = - 0.6, *t* (289) = −8.1, *p* = 1.8 × 10^−14^, R^2^ = 0.2, Robust linear regression, usage captured using the 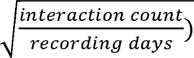.

Across the sample, the smartphone behavior was quantified using the joint interval distribution of next touchscreen interaction intervals (**Fig. 1A**). This distribution consisting of 2500 two-dimensional bins containing the behavioral probabilities was accumulated over a period of 21 days and, as expected, revealed substantial inter-individual differences (**Fig. 1B**) (33). The *behavioral age* model was trained in folds such that on each fold a distinct sub-set of the participants were targeted for predictions (**Fig. 1C)**. The model performed with mean absolute error (MAE) of 6.6 years, and the chronological vs. behavioral age correlated well (spearman *ρ* = 0.8, **Fig. 1D**, see Fig. S3). At the individual level, the distribution of next-interval dynamics was generated daily based on the previous 21 days. The trained *behavioral age* model, applied in folds so as to predict individuals not used in the training, converted the complex distributions into a time series of predictions (**Fig. 1E**).

The population was bimodally distributed (N = 776, with modes at 23 and 64 years of age), allowing us to address the impact of constraining the population-level training data separately to the older mode (> 40 years of age, n = 473) on both the model performance and the timeseries at the individual level. The constrained model performed worse than when using the full age range of the sample (MAE = 5.3 years, *ρ* = 0.6, Fig. S4). At the individual level, the resulting timeseries also showed similar fluctuations as for the full model although the range of the constrained model was more limited (Fig. S5). For subsequent analyses we focused on the unconstrained model.

### Behavioral age tipping points separate basins of attraction

The timeseries of *behavioral age* was analyzed using the Langevin approach to reveal the stability landscape (25, 39). In brief, this approach consists of fitting a so-called Langevin equation to the time series data to capture the deterministic and stochastic parts, and the stability of the one-dimensional dynamics system is described using the effective potential (25). The *behavioral age* effective potential (**Fig. 2A**) contained basins of attraction (stable equilibrium, making valleys in the landscape) separated by tipping points (unstable points making hilltops in the landscape) in the majority of participants (247 of the 280 participants, **Fig. 2B & C**, for subject selection see Fig. S1). The distribution of stable points was broad (**Fig. 2B**), such that individuals > 65 years of chronological age showed stable points even at the low *behavioral age* of ∼40 years (see Fig. S6). The chronological age of people who did show at least one tipping point (median 63 years) vs. people who had no tipping points (median 64 years) was similar. The absence of tipping points was also not linked to the duration of the recording; median recording duration with tipping points 737 days, and without 751 days.

**Figure. 2.**
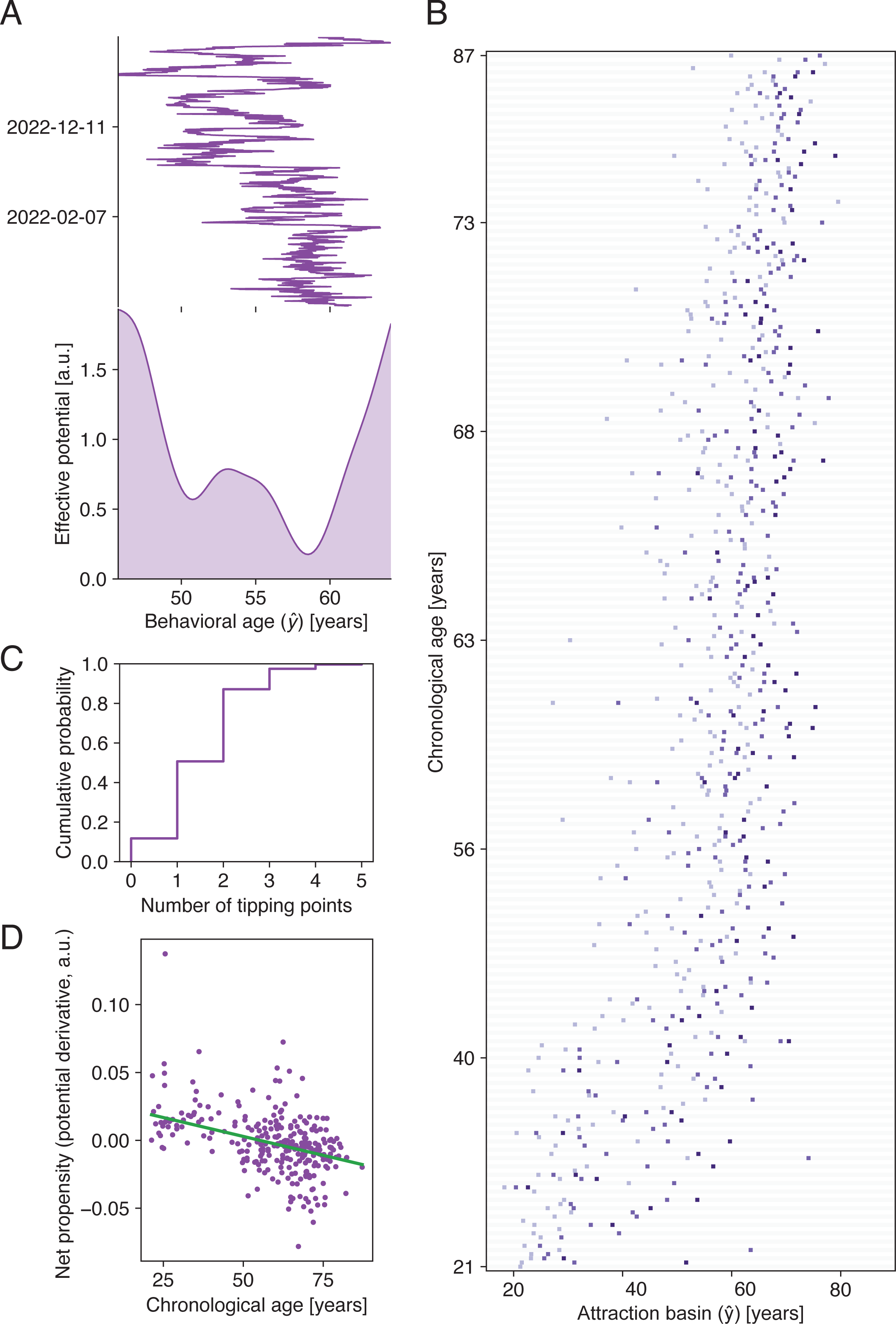
Basins of attraction and tipping points in behavioral age time series. (A) The essence of the fluctuations in *behavioral age* (upper) were captured in an effective potential landscape (lower). Data from one individual to illustrate the idea – here showing two basins of attraction and one tipping point. (B) The location of the basins of attraction across the sample. The locations are sorted by chronological age and within each individual (row) the lighter the shade the *younger* the basin. People with only one basin use the lightest shade. (C) The cumulative distribution of the number of tipping points observed in the population. Most individuals had one tipping point. (D) Adjusted response scatter plot, capturing the relationship between chronological age (the other explanatory variable, *behavioral age* gap, averaged out) and the net propensity of the model fitted dynamical system.

### Stability landscape is unfavorably tilted in people with advanced chronological age

If transitioning to an *older* behavioral state is easier than to a *younger* state, then the effective potential landscape is tilted towards the older state. We captured this in terms of the *net propensity* of the system based on the overall derivative of the landscape (40). Next, we addressed the correlation between this parameter and two explanatory variables: the chronological age and the behavioral *age gap*. The behavioral *age gap* refers to the mean behavioral age model output subtracted by the chronological age (i.e., [*Behavioral age*] – [*Chronological age*] positive Δ is indicative of accelerated aging(32)). The full regression model was significant (*F*(2,273) = 26.1, *p* = 4.1 × 10^−11^) with an *R*^2^ of 0.2 (Multiple linear regression with robust bi-square fitting, n = 3 eliminations, of these due to extreme derivative n = 1, and age gap n = 2). Both the variables were significantly related such that the older the individual, the more negative the derivative (i.e., the landscape tilted towards an older behavioral state, *β*_age_ = - 5.5 × 10^−4^, *t* = −7.0, *p* = 2.0 × 10^−11^, **Fig. 2D**), and the larger the gap, the more negative the derivative (*β* = −7.7 × 10^−4^, *t* = −4.9, *p* = 1.5 × 10^−6^).

### The behavioral resilience indicator alters with chronological age

We focused our analysis on those individuals who had at least one tipping point based on the *behavioral age* landscape (N = 247). We separately analyzed the interindividual differences in the two forms of exit times based on the direction of the exit from the basin of attraction. First, *younger to older*, that is when approaching a tipping point on the older side and second, *older to younger*, that is when approaching a tipping point on the younger side (**Fig. 3A-B**). Pooling the exit times by taking the maximum value revealed a population median exit time of 12 days (inter-quartile range of 38 days), but the maximum exit time was uncorrelated to chronological age and the age gap (*F*(2,235) = 0.5, *p* = 0.6, *R*^2^ = 3.9 × 10^−3^).

**Figure. 3.**
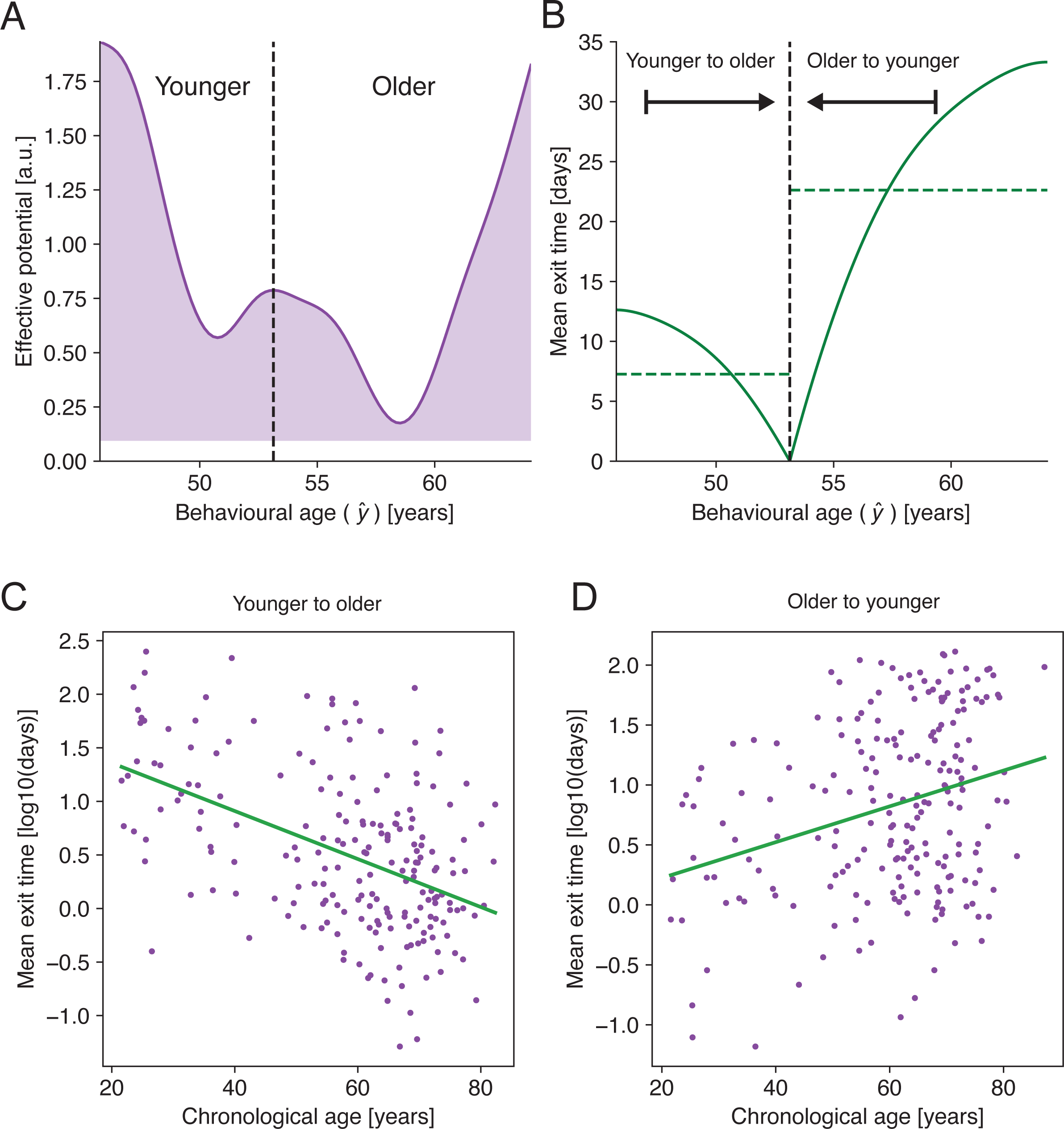
Behavioral Resilience is correlated to chronological age. (A) We focused the analysis of behavioral resilience on those individuals with one tipping point (black dashed line) separating *younger* and *older* basins of attraction. (B) The mean exit time as a function of initial state. Longer exit time indicates a higher behavioral resilience at the corresponding basin. The weighted mean exit time from the alternative basins of attraction towards the boundary (tipping point) are shown in horizontal dashed lines. The exit time from the *younger* basin of attraction (marked as ‘younger to older’) is on the left whereas the exit time from the *older* basin of attraction (marked as ‘older to younger’) is on the right. (C) The *younger to older* exit time diminished with age. (D) The *older to younger* exit time increased with age. Adjusted response scatter plots in C & D, adjusted for the *behavioral age* gap.

We correlated the *younger to older* exit time to the chronological age and the age gap. The full regression model was significant (*F*(2,199) = 20.2, *p* = 1.0 × 10^−8^, *R*^2^ = 0.2), the older the individual the shorter the mean exit time towards the older *behavioral age* tipping point (*β*_age_ = - 2.2 × 10^−2^, *t* = - 6.3, *p* = 1.8 × 10^−09^, **Fig. 3C**, note exit time was normalized using log_10_). And the larger the age gap the shorter the mean exit time (*β* = - 2.6 × 10^−2^, *t* = −3.7, *p* = 2.8 × 10^−04^). For the *old to young* exit time, the full regression model was also significant (*F*(2,208) = 6.4, *p* = 2.1 × 10^−3^, *R*^2^ = 5.7 × 10^−2^), the older the individual, the longer the exit time towards the younger *behavioral age* tipping point (*β*_age_ = 1.5 × 10^−2^, *t* = 3.6, *p* = 4.6 × 10^−4^, **Fig. 3D**). A similar trend was observed for the age gap (*β* = 1.5 _age gap_ × 10^−2^, *t* = 3.6, *p* = 4.6 × 10^−04^). These patterns of results showing age-related differences in the mean exit time were also evident when the analysis was focused on the simple subset of participants with one tipping point separating two basins of attraction (Fig. S7).

### Behavioral alterations underlying the ‘younger’ and ‘older’ basins of attraction

The *behavioral age* fluctuations in some individuals (n = 109) showed just one tipping point separating the younger and the older basins of attraction. We leveraged this relatively simple landscape in conjunction with a feature importance method based on game theory to reveal which two-dimensional joint-interval bins contribute strongly to the two distinct basins (for the feature importance corresponding to the population level model see Fig. S8). At the level of an individual, we estimated the difference in the contributing features between the pairs of two-dimensional bins **(Fig. 4A**). The differences were spread across the joint-interval distribution **(Fig. 4B**). We further separated the two-dimensional bins which contributed inversely (i.e., the higher the behavioral probability, the lower the *behavioral age*) vs. those which contributed directly (i.e., the higher the probability, the higher the *behavioral age*). Taken together, the rapid consecutive intervals contributed inversely such that the higher the probability of behavior the corresponding two-dimensional bins, the lower the *behavioral age* **(Fig. 4C**). The slower interval dynamics appeared to contribute directly such that the higher the probability, the older the *behavioral age***(Fig. 4D**).

**Figure. 4.**
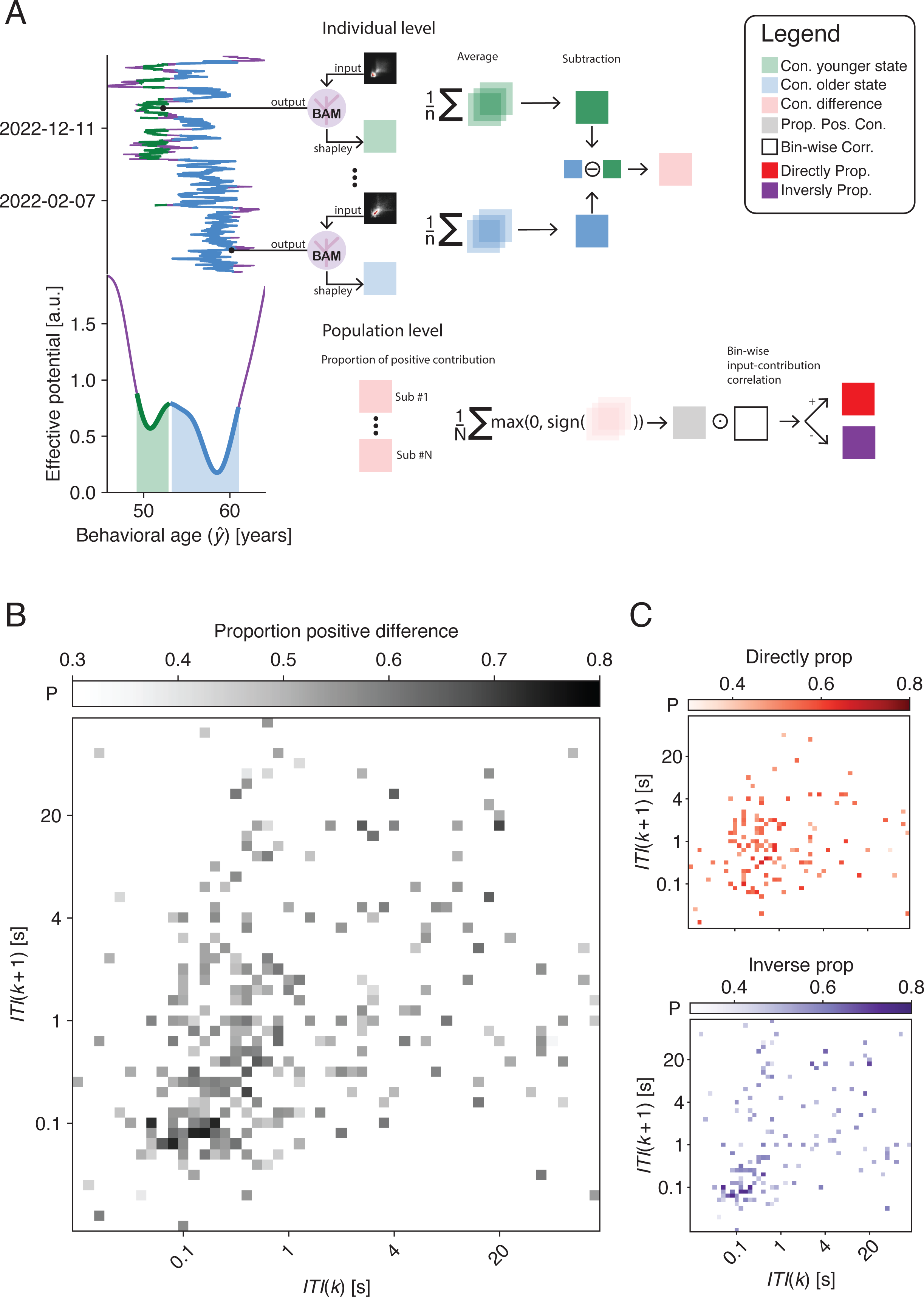
Smartphone behavioral differences underlying the younger vs. older basin of attraction. (A) Schematic view of the analysis used to identify the behavioral features underlying the *younger* vs. *older* basin of attraction. At the level of each individual, Shapley Additive exPlanations was used to obtain maps of feature contribution. We separately considered the contribution maps according to the basin of attraction (*younger* in green, *older* in blue). We averaged the contribution maps in each basin. Next, we subtracted the *older* - *younger* maps to obtain a contribution difference map (‘Con. difference’). At the population level, we use two different descriptors. First, we describe how each two dimensional bin of the JID contributed as in if it contributed more towards the *older* or to the *younger* basin (in grey, ‘Prop. Pos. Con.’). Second, the nature of the change was separated as directly (in red) or inversely proportional (in purple) based on the population wide trend (‘Bin-wise Corr.’). Note in the directly proportional case an increase in the behavioral probability contributes to *older behavioral age* and in the inversely proportional case an increase in the behavior contributes to *younger behavioral age*. (B) The proportion of the population with positive differences in the model contribution. Darker the two dimensional bin the higher the proportion of participants where the behavior contributed more to the *older* basin than the *younger* basin following a change in the behavioral probability density. Conversely, lighter the two-dimensional bin higher the proportion of participants where the behavior contributed more to the *younger* basin following the change. (C) The directionality of the contribution maps. The proportion of the population at a given two-dimensional bin where a positive difference in the model contribution corresponds to directly (red, upper panel) or inversely (purple, lower panel) proportional relationship between the behavior and the behavioral age. For red, upper panel, the darker the two dimensional bin the higher the proportion of participants where the behavior was *older* following an increase in the behavioral probability density. For purple, lower panel, the darker the bin the higher the proportion of participants where the behavior was *younger* following an increase in the behavioral probability. Note, only the feature-selected two-dimensional bins are included in this analysis.

Finally, we checked at the level of each individual whether the basins of attraction and tipping points times could be observed by chance (Fig. S9). For each individual we generated surrogate data by repeatedly shuffling the real time series (1000 bootstrap). In this null distribution any temporal structure is destroyed. In this surrogate data, the number of tipping points and the maximum mean exit time was distinct from what was observed in the real time series. A similar pattern of separation from the null distribution was found when we tested against surrogate data that had the same Fourier spectra as the real data and were generated from a linear Gaussian process (41). For a qualitative perspective on the behavioral age fluctuations see Fig. S10 and the Supporting Information Text.

## Discussion

The rich behavior captured in smartphone interactions varies from person-to-person, and within each individual it varies from day to day. We interpreted the complex behavioral variations in terms of aging. The rich inter-individual behavioral differences could be leveraged by a machine learning model to predict *behavioral age* (32). When the model was applied at the level of each individual, it transformed the rich day-to-day behavioral variations into a time series of *behavioral age* – indicating if the behavior appeared *older* or *younger* from one day to the next. We captured the essence of such short-term fluctuation by using a model tuned to the idea that a complex system operates under the influences of stochastic and deterministic perturbations, where resilience is reflected in the capacity to absorb perturbations without being pushed into an alternative state (25). We find that alternative stable *behavioral age* states are separated by critical tipping points – a property shared with a range of other complex systems such as in ecological systems (16, 19, 25). This allowed us to extend the theory of complex system *resilience* to behavioral aging and reveal how behavioral resilience gradually alters across the adult lifespan.

Individuals abruptly transitioned between *younger* and *older* behavioral states. What does it mean for the behavior to be *younger* or *older*? According to the differences in the feature contributions between these states, the *younger* state was characterized by an increase in the probability of behaviors with fast consecutive intervals (on the scale of ∼100 ms). The *older* state was characterized by a rise in the probability of behaviors involving slow intervals (on the scale of ∼1 s or larger). Taken together with the link between the inter-individual differences in cognitive test performance and the next interval dynamics we speculate that executive functions are weaker in the older state than in the younger state (33). Nevertheless, the alterations in behavioral expression may stem from a complex of intrinsic cognitive abilities and extrinsic factors such as stressful life events. Conventional studies provide limited evidence for day-to-day fluctuations in intrinsic cognitive processes and the variability is marginally reduced in advanced age (42–44). A theoretical framework focused on such day-to-day fluctuations and aging is elusive.

The fluctuations in *behavioral age* can be understood using the theory of tipping points and alternative stable states. At tipping points even small extrinsic or intrinsic forces may cause a cascade of changes to tip the individual from one stable state to another. Our findings support the idea that there are specific periods of openness in aging and in theory there is an opportunity to help the system achieve a more desirable state. In conventional research there is emerging evidence that daily life stressors increase the vulnerability to aging (8, 45). At the tipping point, daily life stressors may drive the behavior toward becoming older, whereas improvements in lifestyle may drive it toward becoming younger. The same forces may have little or no effect in periods when occupying a stable state.

The theory of tipping points and alternative stable states also allowed us to extend an established indicator of resilience in complex systems – mean exit time – to capture the *behavioral resilience*. We observed a broad range of exit times (quartile range between 4 to 40 days). Relatively short mean exit times indicate frequent flipflops between the alternative states, and longer exit times indicate that the individual can more easily absorb perturbations without tipping over. Using this theory, we discovered a vulnerability in the short-term *behavioral age* fluctuations that systematically grows across the adult lifespan. People with advanced chronological age were more vulnerable than the young in tipping over towards an older *behavioral age* and it took them longer to exit towards a younger *behavioral* age. Merging our intraindividual and interindividual observations, as people traverse the decades of the adult life span it becomes gradually more and more difficult to maintain a younger behavioral state day by day or month by month. This idea was further supported by the increased vulnerability linked to the *behavioral age* gap and the net propensity of the system to be more tilted towards the older behavioral states with advanced chronological age.

This perspective of resilience adds to the traditional accounts of age-related resilience in three important ways. First, it uses objective data instead of subjective self-reports and extends a definition of resilience to aging which is mathematically well captured. Second, it is more aligned with the complex realities of behavior by considering the system under continuous intrinsic or extrinsic forces as opposed to well isolated life events. Third, instead of the straightforward traditional account of reduced resilience in the elderly our results offer a more nuanced account where the behavioral resilience is particularly vulnerable to forces that push the system towards an older state but less tolerant of forces that push the system towards a younger state.

It is commonly held that aging constitutes slow irreversible cognitive and behavioral decline spanning decades. Here we extended the theory of alternative stable states and tipping points to rapid age-related behavioral fluctuations in behavior. We find that the essence of the rapid fluctuations gradually alters across the lifespan – thus indicating a bridge between the time scales of days to decades. There is a growing list of isolated perturbations that may alter the pace of aging – from retirement to hospitalization (40, 41) – and identifying such factors and their role is an important function of aging research. Our work indicates the need of another pilar that focuses on stochastic rather than isolated perturbations. These stochastic perturbations (such as linked to busy traffic, spilling wine, forgetting to phone a friend, or missing an appointment), and the behavioral responses to these, help quantify resilience and the structural vulnerabilities in behavioral states. Alternative stable states separated by critical tipping points may be ubiquitous among complex systems – including human aging – and impel the common scientific goal of increasing the resilience of the most desirable states.

## Methods

### Participants

We recruited self-reported healthy participants through the agestudy.nl research platform. This platform leveraged the Dutch Brain Registry of research participants (48). Participants of this platform have been previously reported (31, 33). Furthermore, we pooled archived smartphone data from healthy individuals that contained the self-reported age (37, 49) in terms of the month and year of birth. The health status was confirmed using the RAND SF-36 questionnaire and the mental and physical health components were derived from this score (50, 51). All participants with greater than 10 days of data were included in this study (N = 776, 490 female, 275 male, 11 unreported). The participants provided informed consent which was either written or in the form of a website click (when recruited via agestudy.nl). This study was approved by the local ethics committee (Ethics Commission of the Institute of Psychology, Leiden University).

### Smartphone touchscreen interaction recordings

We recorded smartphone behavior by using a background app introduced (38) and described (31) earlier. Briefly, the users installed the TapCounter app (QuantActions AG, Switzerland) available on the Google Play Store (Google, USA). Each participant was allocated a unique alphanumeric identifier which was entered into TapCounter – the identifier also served to pool data from multiple smartphones belonging to the same user. The app captured the timestamp (in UTC milliseconds) of touchscreen interaction conducted by the user across all apps on the smartphone in addition to the label of the foreground app in use (such as *com.facebook.katana*) and low resolution (city/town level) location information. The screen switch *ON* and *OFF* events were also gathered by the app. All of the data was streamed from the app to a cloud platform tethered to the unique identifier (Taps.ai, QuantActions AG). The data was parsed using a raw data parser provided by QuantActions. During the recording period the collection was routinely (weekly) monitored online using taps.ai, and in the event of any data discontinuity the user was contacted to resume the data collection as noted earlier (31).

### Behavioral age model based on smartphone touchscreen interactions

A decision tree regression model was trained to estimate the chronological age based on the smartphone touchscreen interactions – we termed these predictions as *behavioral age* (32). Briefly, the next-interval dynamics of smartphone touchscreen interactions was captured using a joint interval distribution (JID) (30). This distribution captures the probability of observing interaction triads of various next-interval dynamics in 2500 two-dimensional log_10_-spaced bins spanning from ∼30 ms to ∼100 s (Fig. 1A). Note, only within-session (the period between screen *ON* and *OFF* events) intervals were considered towards the JID. The accumulation period for the JID was 21 days. A JID was extracted every 60 days in participants with data spanning longer than 80 days. The JIDs were separated in 10 folds for cross validation training and testing.

The model training procedure used here consisted of a feature selection step (which was absent in the original smartphone age estimating model (32)). The features consisted of the 2500 two-dimensional bins of the JID and three additional features calculated across the period of accumulation: the median inter-touch interval, the interquartile range and the median number of taps per day. From these 2503 features, 250 features were selected using an iterative method that reduced the number of features based on the least significant features at each iteration using the Gini importance contribution method for XGBoost (52). The naïve model was trained using standard parameters (see code availability section).

We iteratively established the XGBoost parameters using grid search of 1-2 parameters on each run. On each run the best values were selected based on the average R^2^ across a 10-fold cross validation from the training set. The cross-validation with basic parameters was used to form an initial guess of the number of boosting rounds needed. The grid search parameters are shared on the code repository (see below). A final cross-validation was performed with the searched optimal parameters and a learning rate of 0.01 to establish the optimal number of boosting rounds with an early stopping of 5 rounds to avoid overfitting.

### Behavioral age timeseries

The trained behavioral model was applied in folds on each *individual* such that the trajectories were based on the individuals not used towards the training. For each of the individuals with at least one year of data (N = 318), daily JIDs (accumulating 21 days of smartphone behavior) were used as the model input resulting in one behavioral age prediction per day. We next eliminated individuals with more than 7 continuous days without predictions, or exceeding a threshold (50%) of missing predictions (say due to insufficient number of touches). This left N = 291 individuals for the Langevin fitting procedure to estimate the basins of attraction and tipping points.

### Estimating basins of attraction and tipping points from behavioral age timeseries

We fitted a Langevin equation to the timeseries to describe the state variable (*x*):

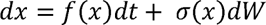

The *f(x)* part of the function is the deterministic part, *σ(x)dw* captures the increments of a Weiner process with uncorrelated random fluctuations and its standard deviation linked to the state. The fitting procedure is described in detail elsewhere (39). Briefly, the estimation of drift and diffusion functions used the covariance matrices of these quantities. These estimates were obtained with kernel-weighted averages in two steps, first using the difference and then the square of first difference of the timeseries *(x)*. The Gaussian kernel was defined by evenly dividing the range of values of the timeseries according to a mesh and a bandwidth. The interplay between these two parameters will result in a more or less smooth estimate of these functions. The details of these operations can be found in (39). While we attempted other methods to estimate drift and diffusion (24), the *mesh* method as indicated above yielded the smoothest results even with relatively short timeseries. This is due to the possibility of choosing the level of smoothness by varying the bandwidth of the kernel and the density of the mesh. We set the density of the mesh to: 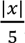 b and width to: 0.3 × σ(*x*), according to (39). This analysis failed (yielding complex coefficients) in 10 individuals yielding in a final N of 281 individuals.

### Estimating net propensity and the mean exit time

Towards net propensity, we estimated the derivative of the effective potential landscape as reasoned here (40). The mean exit time was estimated as reasoned here (25). The exit time from a left basin of attraction to the right tipping point (*younger to older*) was separately considered from the exit time from the right basin to the left tipping point (*old to young*). As in the prior work, exit times in between two or more tipping points were not considered.

### Surrogate data for null distributions

We generated surrogate behavioral age timeseries for all the individuals with at least two basins of attractions using the following two of the methodologies described in (41). The first method involved randomization, towards this we resampled the real time series with replacement and created 1000 surrogate time series. This was to test the simple case of whether there is any temporal structure in the data. The second method involved preserving in the surrogate the Fourier spectrum of the real time series, which in turn preserved the autocorrelation properties of the real time series. Towards this we employed a phase randomization process where for each surrogate times series we generated a random phase vectors which was then applied to the real time series’ Fourier spectrum. This new spectrum was then inverted into the time domain to obtain the surrogate time series, this process was repeated 1000 times. This creates a null distribution to test the case whether there is any non-linear structure in the data. The Langevin equation was fitted to the surrogate data and the null distribution was composed of the maximum mean exit time observed from landscapes with the same number of tipping points as in the real data or else the exit time was set to zero.

### Feature importance analysis

To perform the feature importance analysis underlying the distinct basins of attraction we selected individuals who showed 2 basins of attraction separated by a tipping point (N = 91).To establish the importance of input features underlying the behavioral age fluctuations in each of these individuals we used SHAP (Shapley Additive exPlanations). This is an exact method to calculate feature importance based on the game theoretic approach of Shapely values which captures how much an input feature contributes to the model output (53). That is, a feature with a high contribution strongly influences the output value. In particular, a feature with high contribution is a two-dimensional bin of the JID that is frequently occupied (i.e., high probability density) or infrequently present (i.e., low probability density), and responsible for placing the individual in a particular *behavioral age*. We separated the time series of Shapely values into two bins corresponding to the two basins of attraction – the *older* and *younger* basin. The younger bin held all the points in the time series that fell in the left basin of attraction (< tipping point). Conversely the older bin held all the points in the time series that fell in the right basin of attraction (> tipping point). We averaged the Shapley Values resulting in 250 features large contribution maps corresponding to each basin (note, feature selection from the 2500 JID features in the behavioral age model). Next, we subtracted the older average – younger average to obtain the contribution difference at the level of each individual.

To obtain a population level overview from the contribution differences, we focused on the direction of the difference. That is each of the 250 two-dimensional bins were either: +1, higher contribution at the older basin, or-1, lower contribution in the older basin, and the map was binarized marking the bins that contributed positively. The values were averaged across the population resulting in a contribution where the value between 0 to 1 indicated the ratio of participants contributing positively at the given two-dimensional bin. This population overview contained two forms of contributions: one, where a higher probability density resulted in higher behavioral age prediction (directly proportional) and two, where a lower probability density resulted in higher prediction (inversely proportional). To separate these to different types, we used a population-level classification. Towards this, we estimate the correlation type across the sample, and next multiplied this to separate the direct or inverse proportion contributions. The underlying equations are annotated in the Supporting Information Text (along with the steps used towards Fig. S8).

### Linear regression analysis

We performed robust linear regression analysis to link the inter-individual differences in the net propensity and the mean exit times, to the chronological age and the *behavioral age* gap:

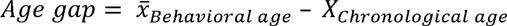

In total we ran 5 different robust linear regression analyses (MATLAB, Mathworks, Natick). We express the *p* values in the text and Bonferroni corrected relationships (α = 0.05, α_Bonferroni_ = 0.01) are depicted using solid lines in the figures.

## Data Availability

The timeseries of age predictions are shared on dataverse.nl upon publication.

## Code Availability

The codes used to process the timeseries of age predictions are shared on: https://github.com/codelableidenvelux/ML_Age_Trajectory_2023

## Funding and acknowledgment

This study was funded by Velux Stiftung (project no. 1283, awarded to AG with KRR as co-applicant). EC was supported by the SNSF Early Postdoc.Mobility (no. 199692 awarded to EC with AG as host). The authors would like to thank the student and staff researchers behind the data collection platform agestudy.nl. We thank Sander Nieuwenhuis for his help in editing this manuscript and improving its readability.

## Author contributions

AG conceived the study aided by EC. EC processed the data aided by AG. AG drafted the report aided by EC. RR edited the manuscript.

## Data and code sharing

The time series of age predictions shall be shared on dataverse.nl and the codes to analyze the data are shared on GitHub.

