## Supplementary figures and images for "Age-related behavioral resilience in smartphone touchscreen interaction dynamics"

### Fig. S1

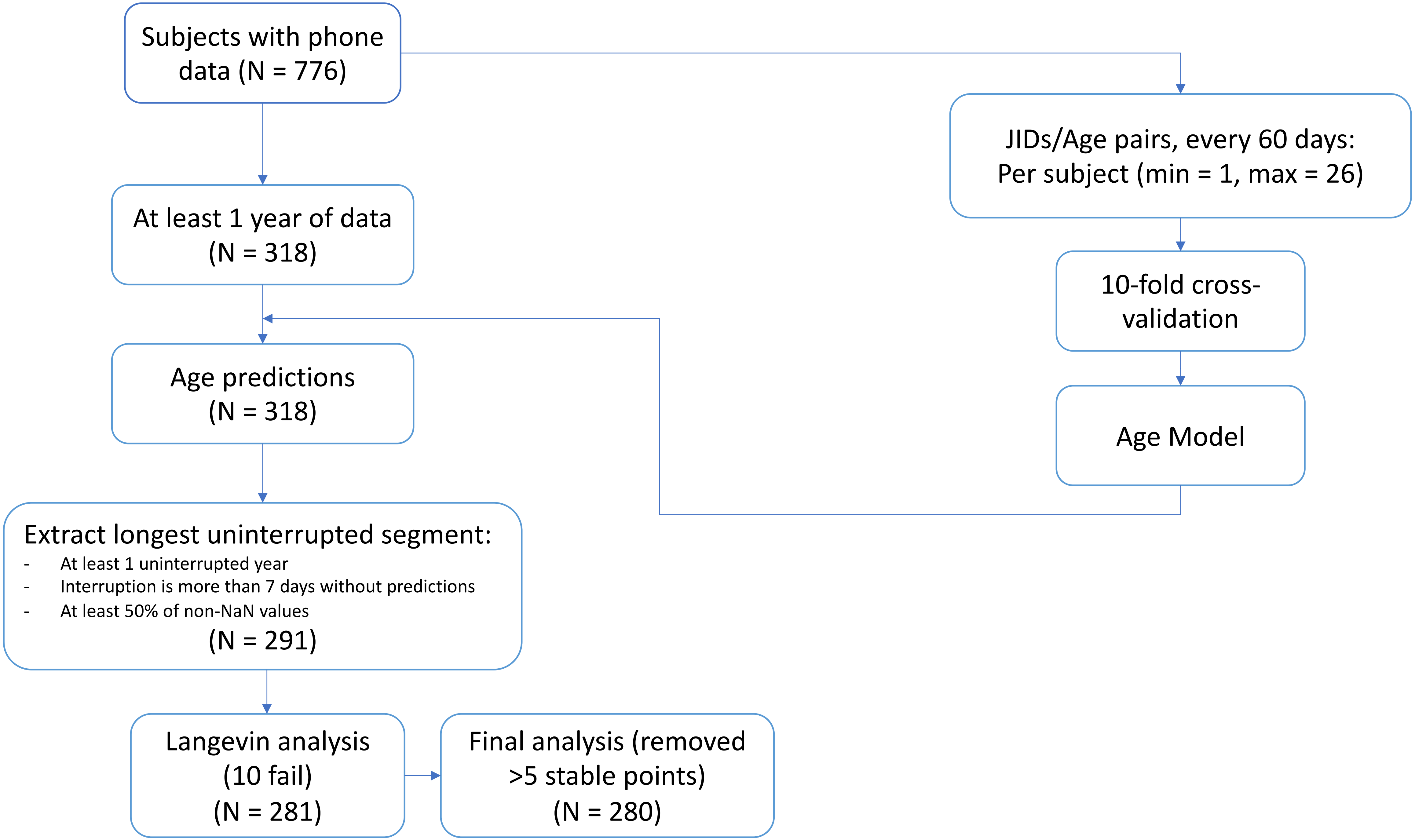

### Fig. S2

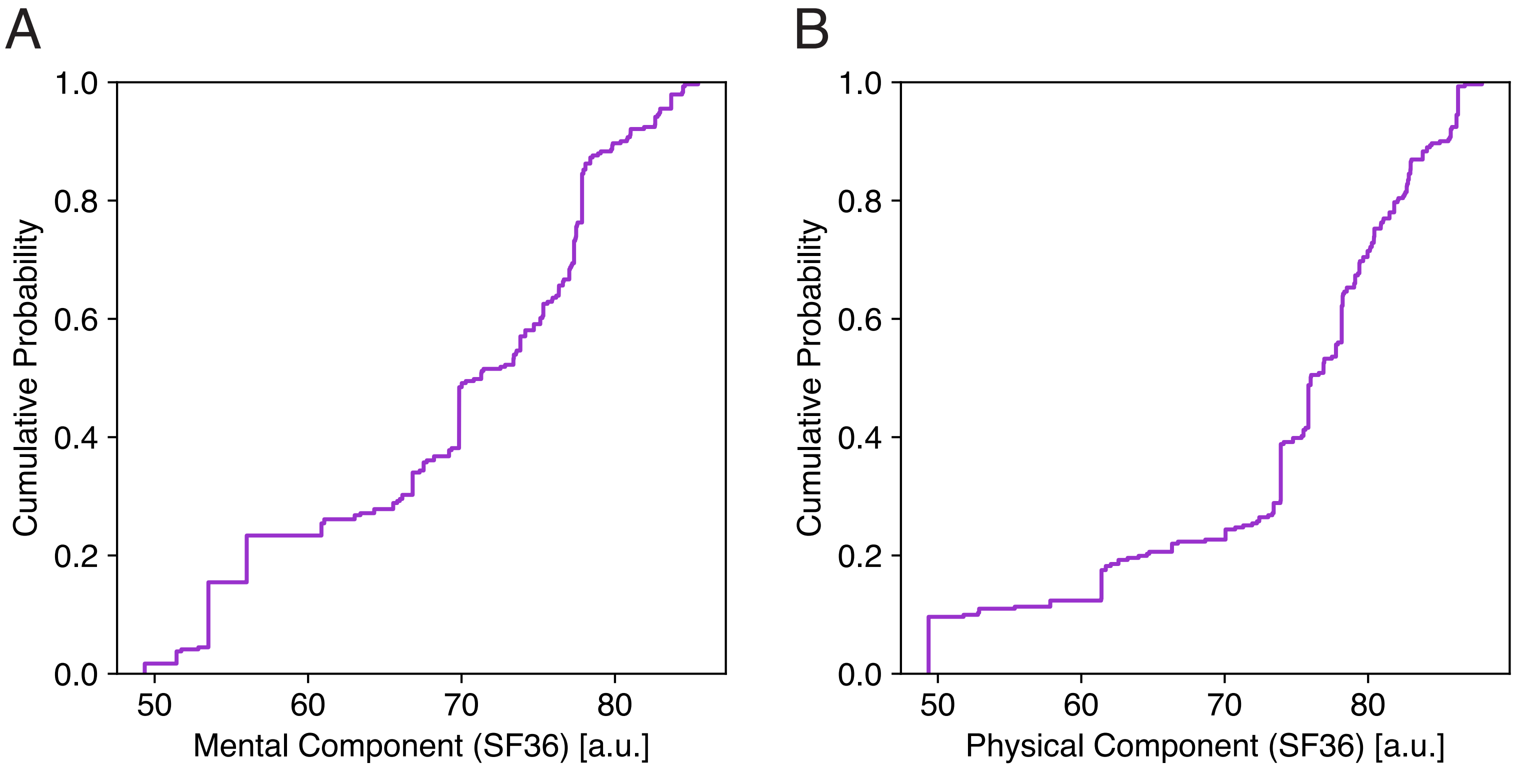

### Fig. S3

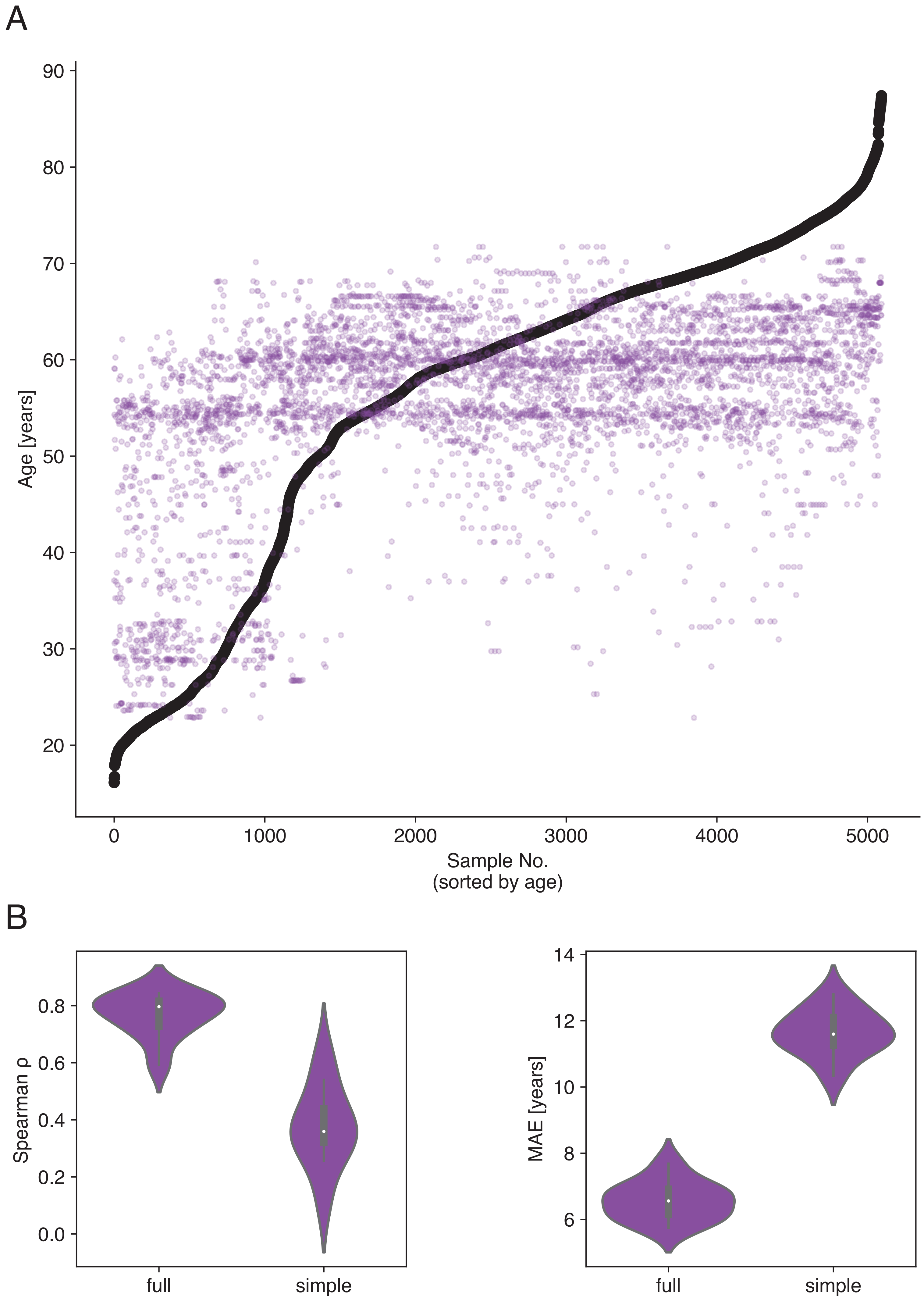

### Fig. S4

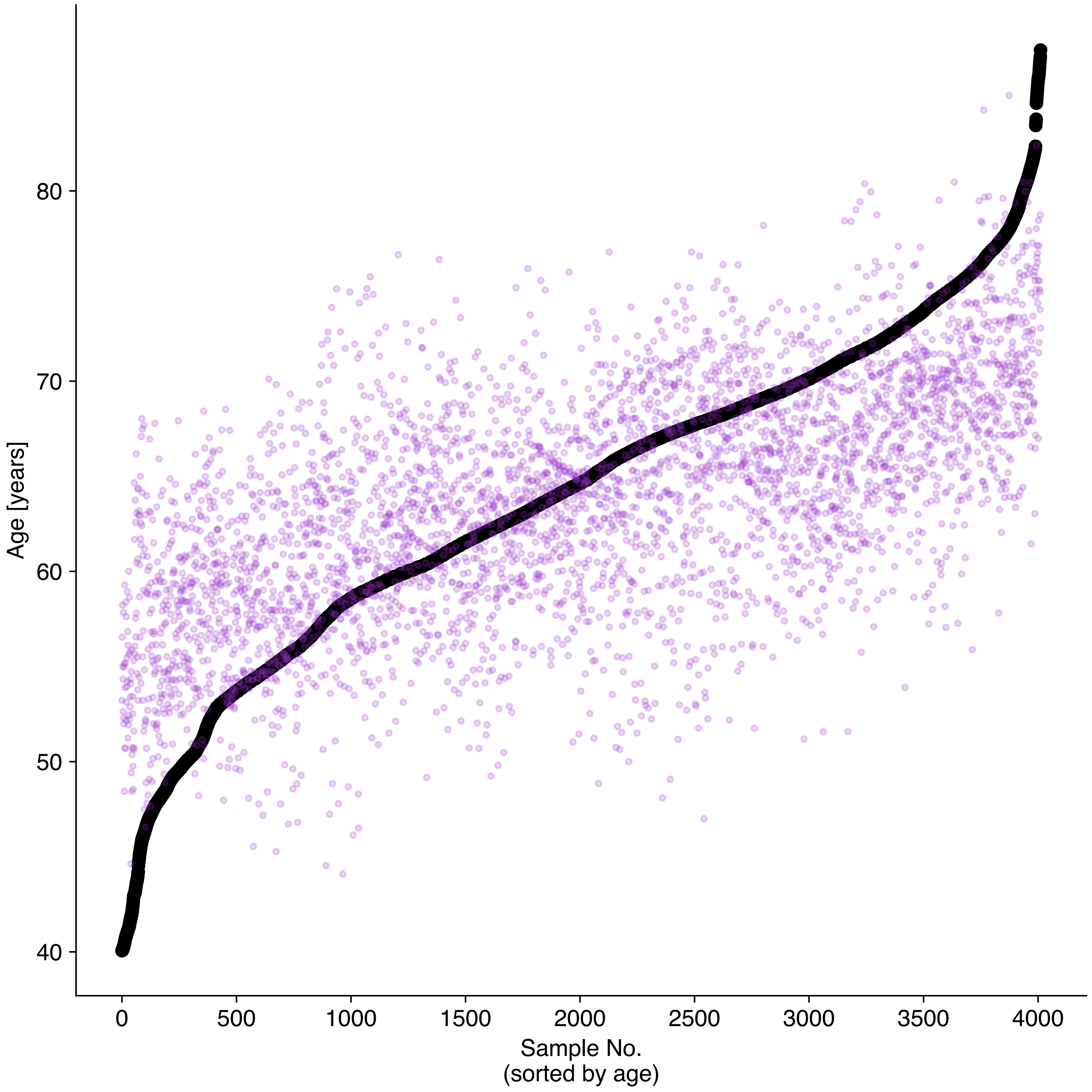

### Fig. S5

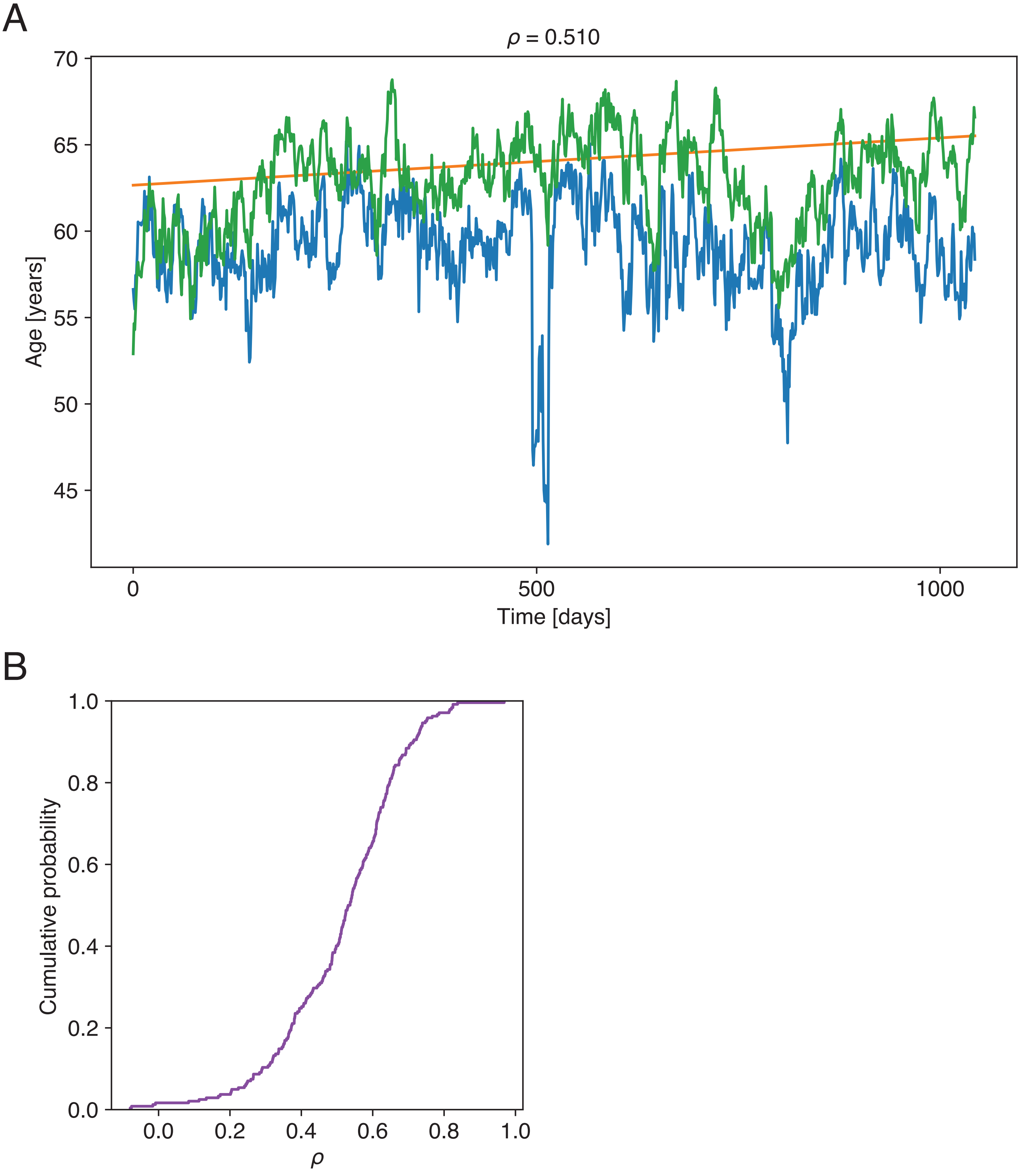

### Fig. S6

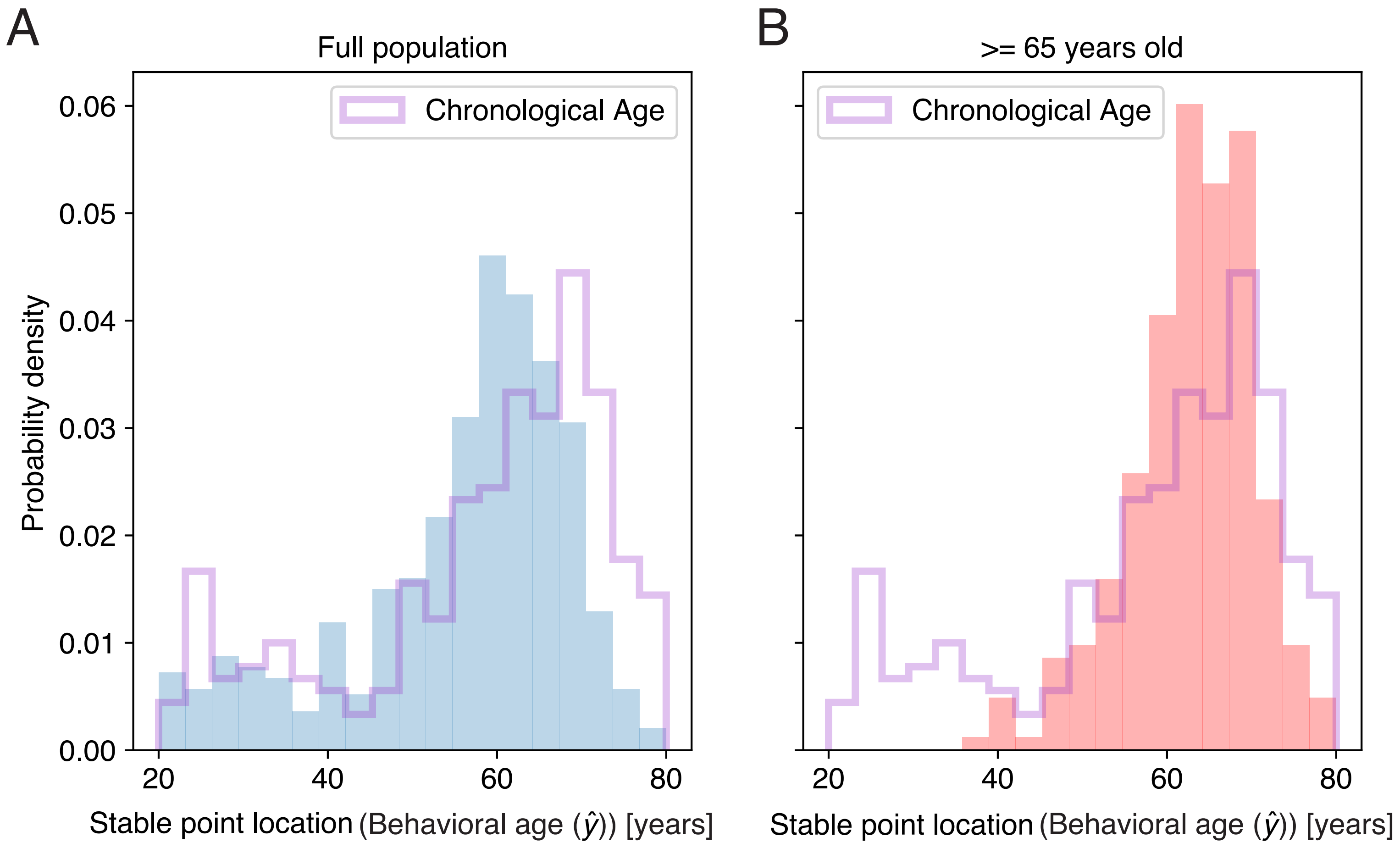

### Fig. S7

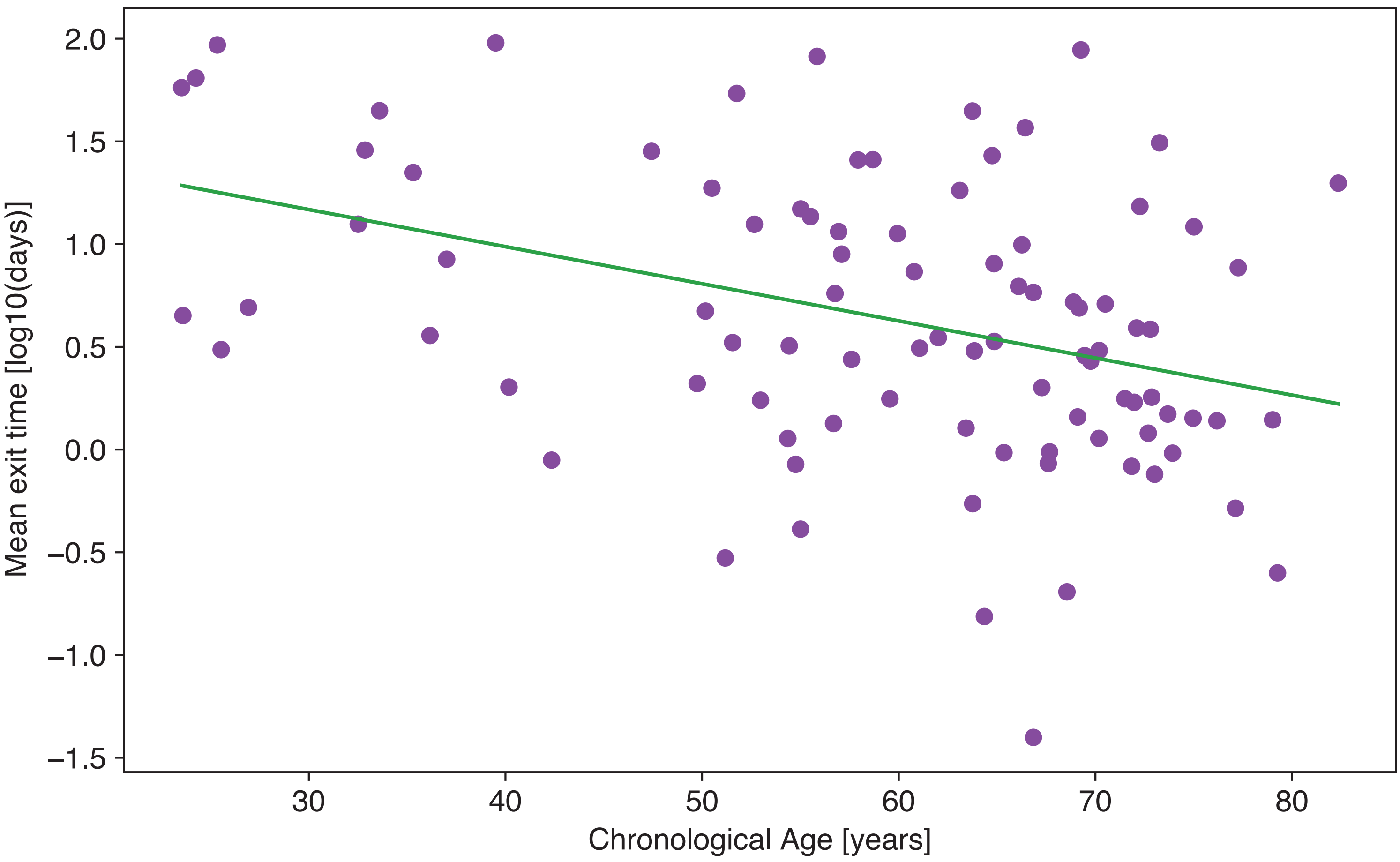

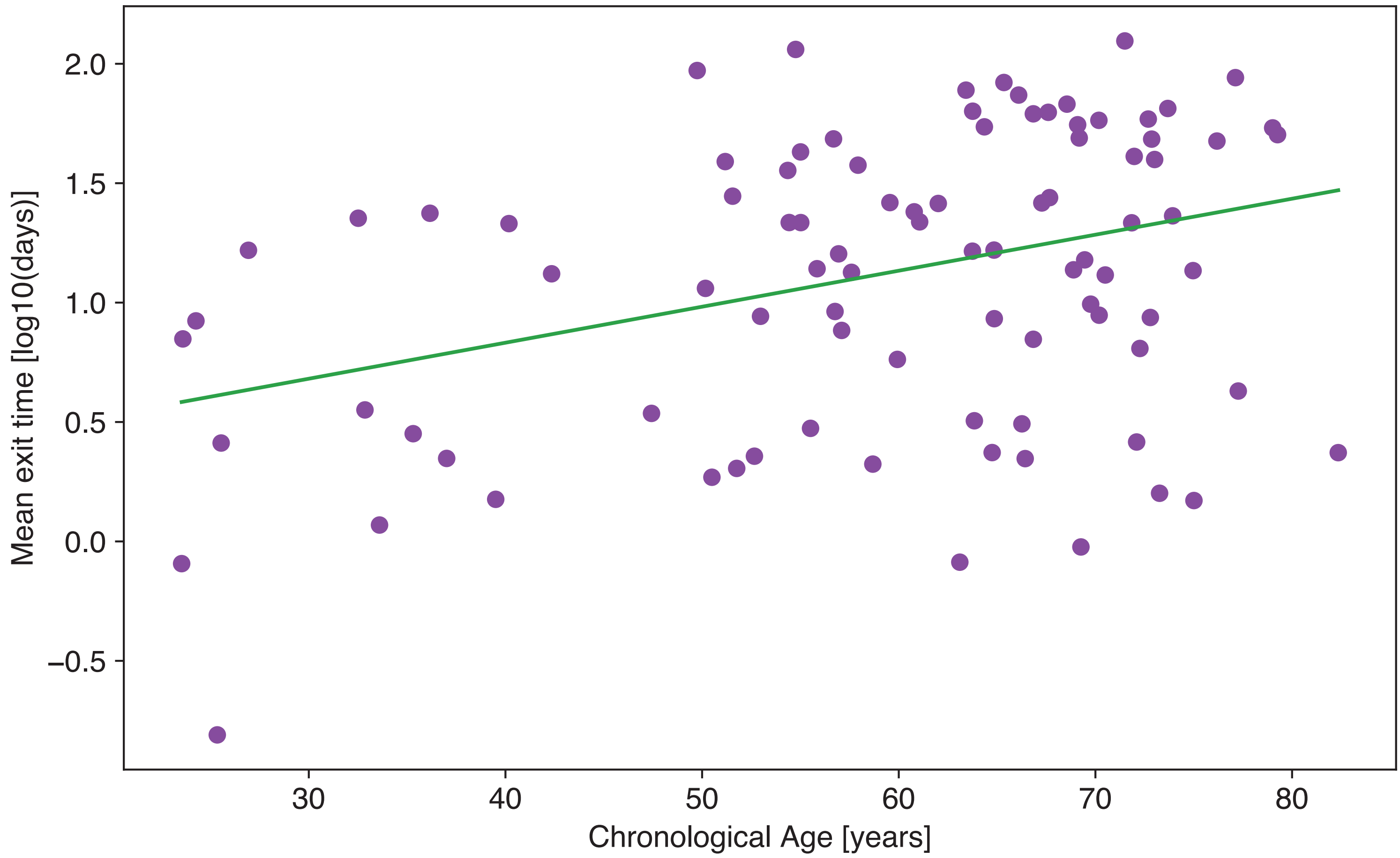

### Fig. S8

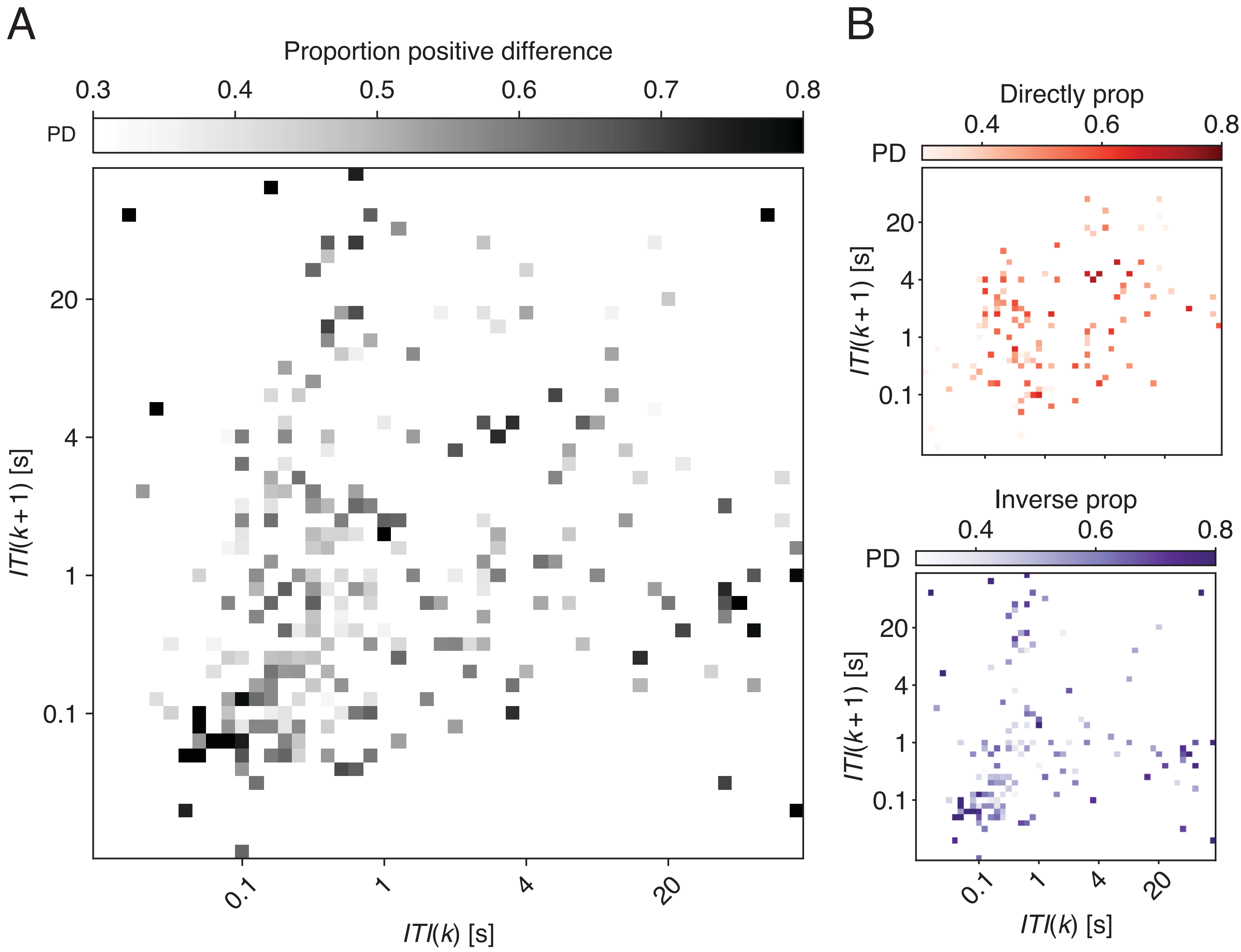

### Fig. S9

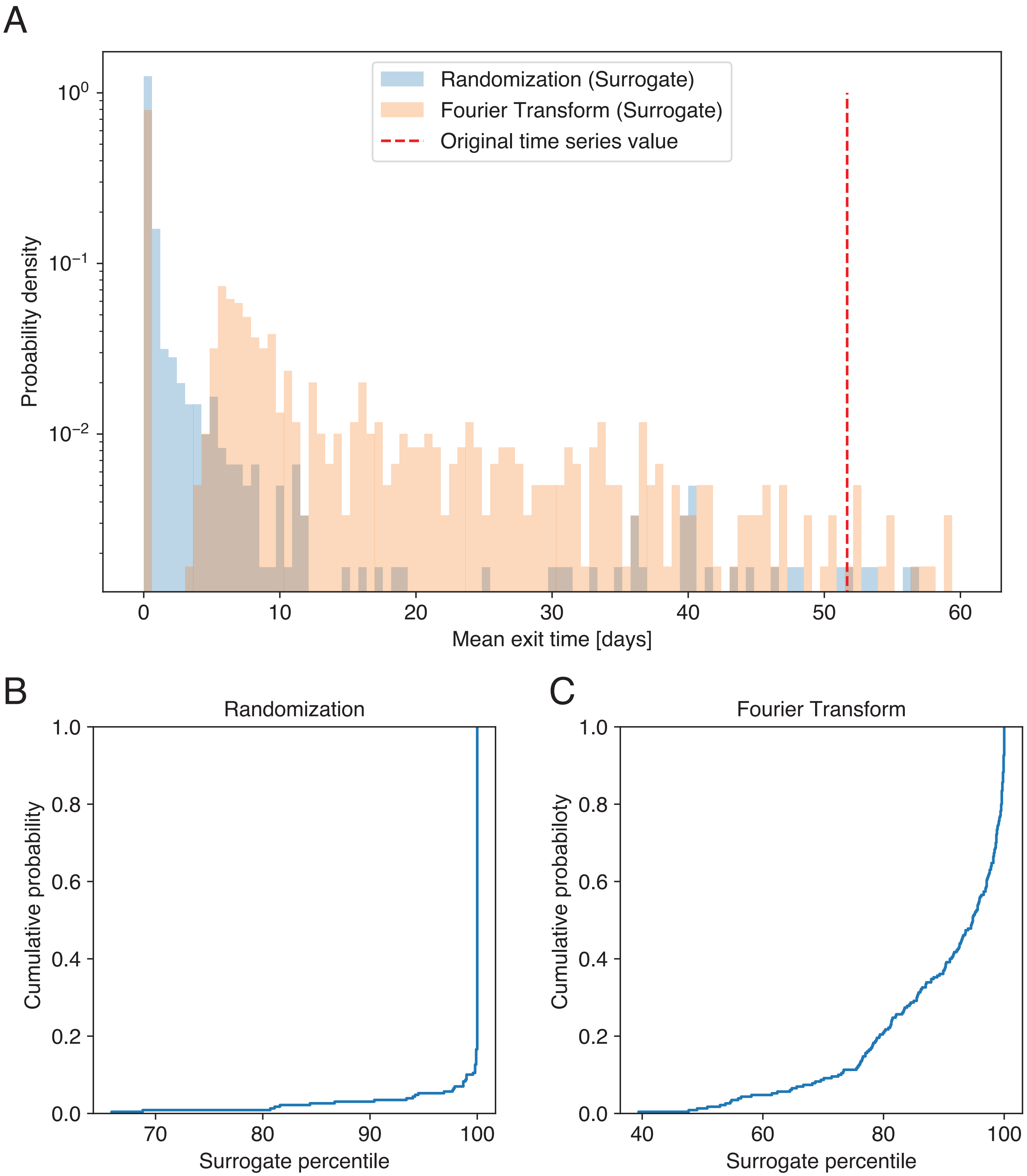

### Fig. S10

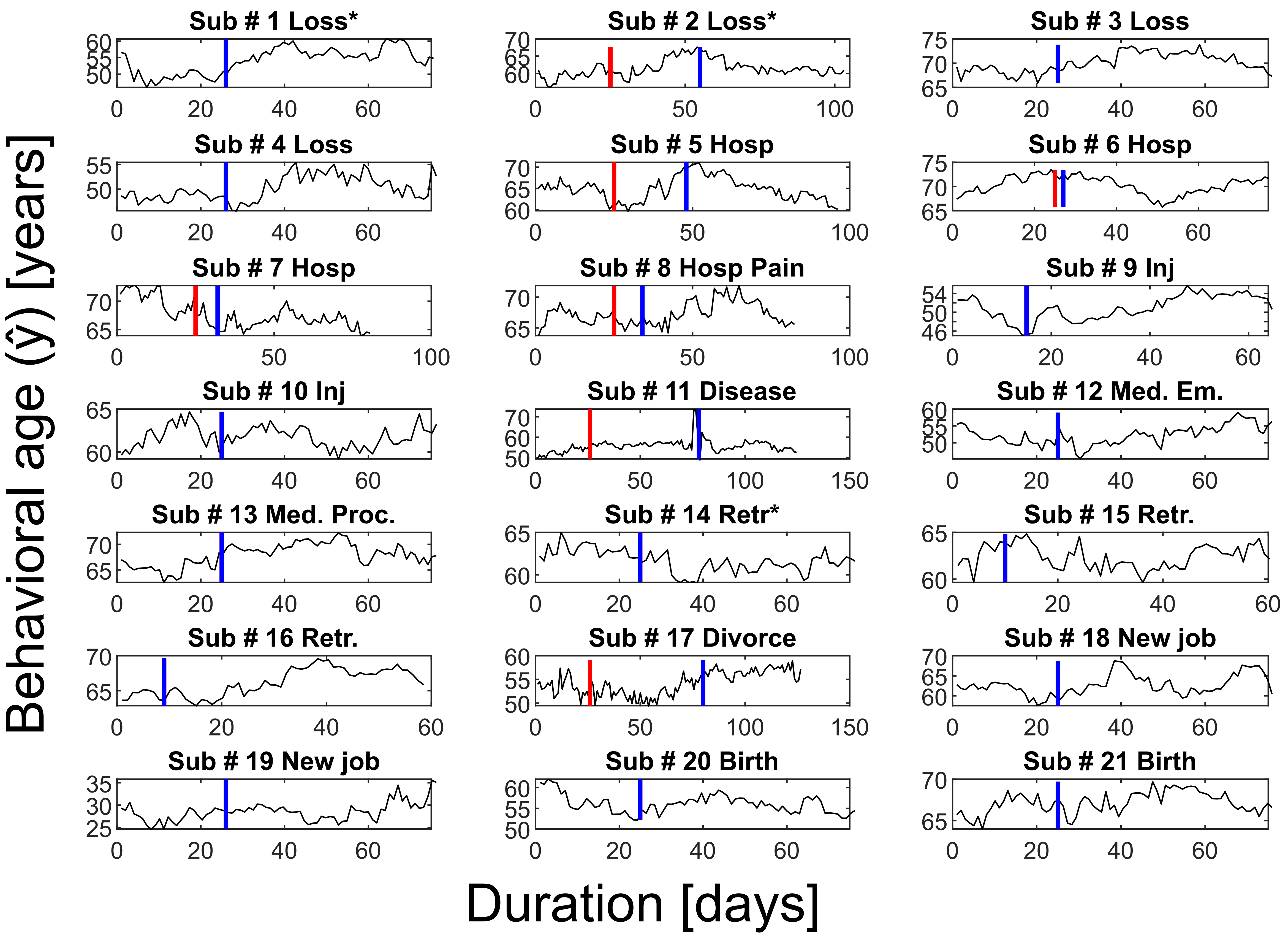
